# Cell-type specific EWAS identifies genes involved in HIV pathogenesis and oncogenesis among people with HIV infection

**DOI:** 10.1101/2023.03.21.533691

**Authors:** Xinyu Zhang, Ying Hu, Ral E. Vandenhoudt, Chunhua Yan, Vincent C Marconi, Mardge H. Cohen, Amy C Justice, Bradley E Aouizerat, Ke Xu

**Author notes:** Corresponding to Ke Xu, MD, PhD Bradley E Aouizerat, Ph.D.

## Abstract

Epigenome-wide association studies (EWAS) of heterogenous blood cells have identified CpG sites associated with chronic HIV infection, which offer limited knowledge of cell-type specific methylation patterns associated with HIV infection. Applying a computational deconvolution method validated by capture bisulfite DNA methylation sequencing, we conducted a cell type-based EWAS and identified differentially methylated CpG sites specific for chronic HIV infection among five immune cell types in blood: CD4+ T-cells, CD8+ T-cells, B cells, Natural Killer (NK) cells, and monocytes in two independent cohorts (N_total_=1,134). Differentially methylated CpG sites for HIV-infection were highly concordant between the two cohorts. Cell-type level meta-EWAS revealed distinct patterns of HIV-associated differential CpG methylation, where 67% of CpG sites were unique to individual cell types (false discovery rate, FDR <0.05). CD4+ T-cells had the largest number of HIV-associated CpG sites (N=1,472) compared to any other cell type. Genes harboring statistically significant CpG sites are involved in immunity and HIV pathogenesis (e.g. *CX3CR1* in CD4+ T-cells, *CCR7* in B cells, *IL12R* in NK cells, *LCK* in monocytes). More importantly, HIV-associated CpG sites were overrepresented for hallmark genes involved in cancer pathology (*FDR*<0.05) (e.g. *BCL family, PRDM16, PDCD1LGD, ESR1, DNMT3A, NOTCH2*). HIV-associated CpG sites were enriched among genes involved in HIV pathogenesis and oncogenesis such as Kras-signaling, interferon-α and −γ, TNF-α, inflammatory, and apoptotic pathways. Our findings are novel, uncovering cell-type specific modifications in the host epigenome for people with HIV that contribute to the growing body of evidence regarding pathogen-induced epigenetic oncogenicity, specifically on HIV and its comorbidity with cancers.

## Introduction

With successful antiretroviral therapy (ART), people with HIV (PWH) have a similar lifespan to the general population [1]. However, the healthspan of PWH remains 9 years shorter [1] because of a high burden of comorbid chronic diseases such as cardiovascular diseases [2], diabetes, and non-AIDS-related cancers [3]. The prevalence of non-AIDS-related cancers among PWH is significantly higher compared to that among the people without HIV (PWoH) [4, 5], especially in PWH who are not virally suppressed [6, 7]. Elevated cancer incidence may be due to the high prevalence of cancer risk factors such as substance use and other co-infections [8]. It is important to characterize the underlying mechanisms involved in HIV pathogenesis that may contribute to cancer emergence for PWH.

Host epigenetic modifications play critical roles in HIV-1 induced cellular reprogramming at different stages of HIV-1 pathogenesis, including viral integration, maintenance, activation, or silencing [9]. Upon the integration of HIV-1 into the host genome, chromatin in the infected cells undergoes profound reorganization to control the virus by affecting proviral long terminal repeat (LTR) promoter complex formation [10]. HIV-1 proteins (e.g. Tat) in turn change the cellular environment to facilitate virus survival and replication by disrupting chromatin structure and altering gene expression in the host cell. For example, expression of histone methyltransferase (*DNMT3A*, *DNMT3B*) and histone deacetylase (*HDAC2* and *HDAC3*) genes are significantly upregulated in host cells infected with HIV-1 [11]. Additionally, although some cells infected with HIV have a normal expression of the BCL family of anti-apoptotic proteins permitting apoptosis and viral propagation, other infected cells overexpress these proteins [12], thereby increasing the risk of cancer and promoting persistence of latently infected cells [13], which poses a barrier to HIV eradication. Epigenetic reprogramming in immune cells persists in chronically infected cells. Such dynamic virus-host genomic interaction results in distinct epigenetic profiles among different immune cell types in response to environmental changes. A common hallmark of pathogen-induced accumulation of DNA methylation maladaptation in multiple genes is an increase in risk for oncogenesis and cancer, which is estimated to occur for as much as 20% of cancers [14]. One example is hepatitis B virus-induced DNA methylation alteration of tumor-suppressor genes *p16*, *p21*, *CDH1*, and *SOCS1* that contributes to hepatocellular carcinoma [15]. However, the role of HIV-associated epigenetic alterations in carcinogenesis has not been explored.

Epigenome-wide association studies (EWAS) of the host methylome have identified numerous significant CpG sites for different stages of HIV infection. During the acute stage of HIV infection, up to 22,697 methylation sites are altered by HIV-1 [16]. ART initiation reverses patterns of DNA methylation in less than 1% of the altered CpG sites, leaving the majority of CpG sites with methylation states persisting into the chronic stage even among virally suppressed individuals [16]. Independently replicated studies have identified several CpG sites and genes associated with HIV infection, including three *NLRC5* promoter CpG sites that are less methylated in PWH with or without ART compared to PWoH [17–20]. DNA methylation profiles are predictive of HIV progression, frailty, and mortality for PWH [21–23]. These findings demonstrate the importance of methylation mechanisms in HIV infection and HIV-related comorbidities.

While EWAS in whole blood have identified CpG sites such as *NLRC5* that have been replicated by different studies, they provide limited insight into the epigenetic modifications of specific immune cell populations that underlie the pathogenic effects of HIV. HIV-1-induced alterations to the epigenomes of specific immune cell types remain unknown. Immune cells that originate from different lineages show distinct DNA methylation patterns [24, 25]. Thus, conventional blood-or peripheral blood mononuclear cell (PBMC)-based EWAS are likely to confound cell-type specific CpG sites for HIV-1 infection. EWAS signals identified from heterogeneous cells are likely to result from consistent CpG methylation signals across cell types or a strong cell-type specific signal that exceeds noise from inconsistent methylation of the same CpG in other cell types. However, only a few studies have conducted EWAS in specific cell types to identify HIV-1 associated CpG sites, and then only in a few cell types and in limited sample sizes. For example, one study showed that the number of CpG sites changed by HIV-1 were approximately 100 times greater in monocytes than CD4+ T-cells in the acute stage of HIV infection [16]. Little is known about the impact of HIV-1 infection on the epigenome of other cell types such as B cells, CD8+ T-cells, natural killer (NK) cells, in addition to monocytes and CD4+ T-cells. DNA methylation in specific cell types plays an important role in responding to, but is ultimately altered by, HIV-1 infection [26, 27].

Cell-type specific DNA methylome profiling of large population samples is technically challenging and cost prohibitive, especially for less abundant cell types in blood. Recently, computational methods have been developed and applied to deconvolute cell-type specific methylation signals from bulk PBMCs or whole blood samples [28–31]. You et al. (2021) successfully dissected several well-known smoking-associated hypomethylation signatures that derived from myeloid lineage immune cells [32].

In this study, we hypothesized that alterations in CpG methylation in the genome of individuals with chronic HIV differ among immune cell types and that HIV-associated CpGs are enriched among genes involved in HIV pathogenesis, but potentially other comorbid conditions including cancer development and progression. We applied a tensor composition analysis (TCA) method to computationally deconvolute cell-type specific methylation data in PBMCs without sorting cells [33]. We first validated the TCA deconvoluted methylomes by direct bisulfite DNA sequencing of sorted CD4+ T-cells, CD8+ T-cells, and monocytes from the same PBMC specimen for a subset of the sample (N=29). We then deconvoluted methylation data from whole blood or PBMCs into CD4+ T-cells, CD8+ T-cells, B cells, NK cells, and monocytes in 1,134 samples. These cell-type specific EWAS for chronic HIV infection were conducted in two cohorts: the Veteran Aging Cohort Study (VACS)[34] for men (N=702) and the Women’s Interagency HIV Study (WIHS)[35] for women (N=432) **(Supplemental sTable 1)**. To the best of our knowledge, this is the first and largest cell-type specific EWAS for chronically infected PWH in a predominantly African American population. A flowchart of analytical strategies is presented in **Figure 1**.

**Figure 1.**
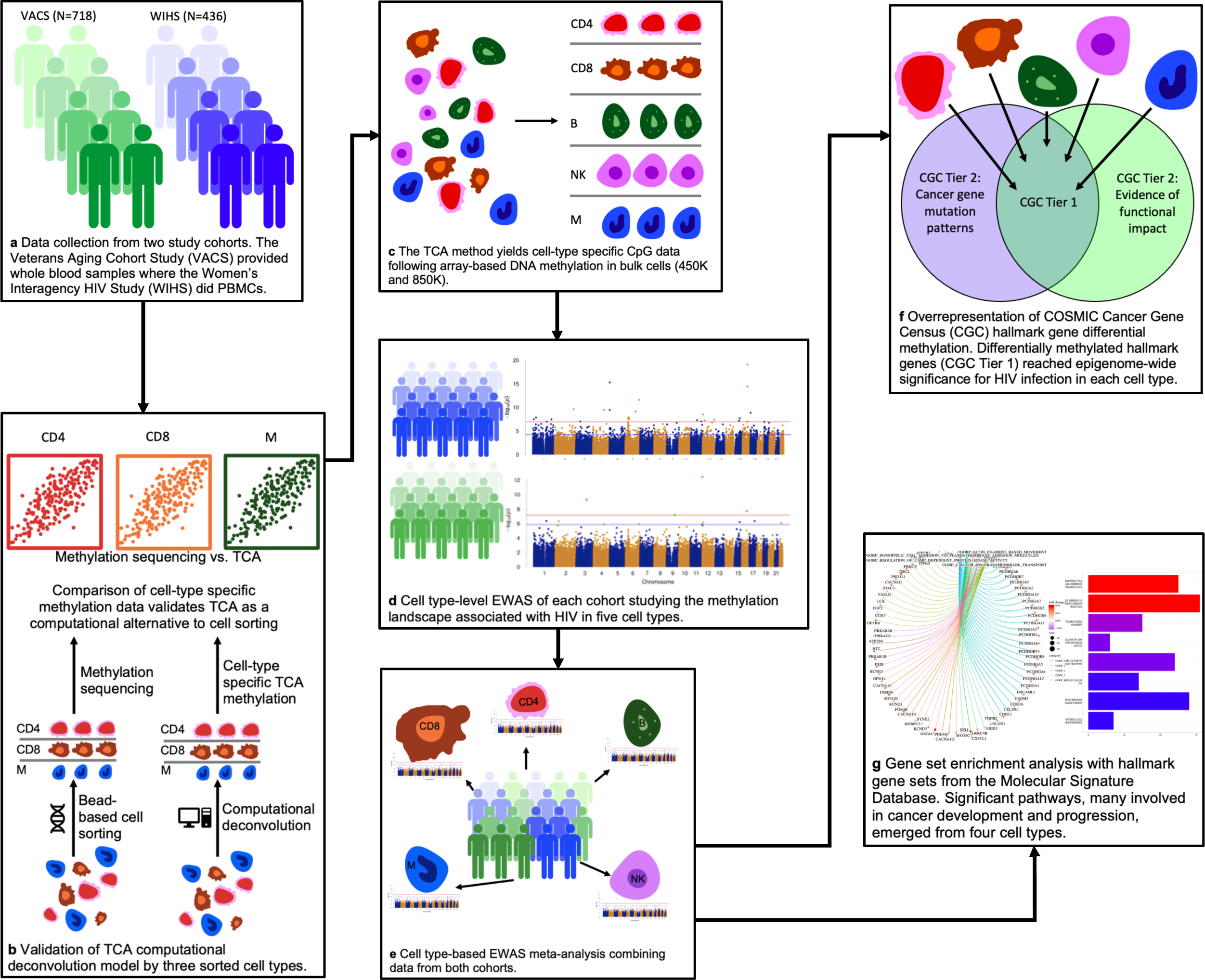
Flowchart of analytical strategies. VACS: Veteran Aging Cohort Study; WIHS: Women’s Interagency HIV Study; TCA: Tensor Component Analysis; EWAS: Epigenome-wide Association Study.

## Results

### Validation of deconvoluted DNA methylome data measured using capture sequencing in CD4+ T-cells, CD8+ T-cells, and monocytes

We first sought to validate the performance of TCA by comparing the DNA methylomes of CD4+ T-cells, CD8+ T-cells, and monocytes isolated from PBMCs to TCA-deconvoluted DNA methylation data collected employing capture sequencing in a subset of WIHS samples. The methylation value of each CpG site from the bulk (PBMC) sequencing data was deconvoluted to CpG methylation originating from CD4+ T-cells, CD8+ T-cells, and monocytes using TCA. In tandem, a separate aliquot of the same PBMC sample was subjected to cell sorting using magnetic beads and the CD4+ T-cells, CD8+ T-cells, and monocytes obtained were individually subjected to methylation capture sequencing (MC-seq) (**Supplemental Material**).

We compared the methylation β-value for the top 10,000 most-variable CpG sites between the TCA-deconvoluted and the directly measured methylation β-value for CD4+ T-cells, CD8+ T-cells, and CD14+ monocytes. We found a high correlation of methylation values between the two approaches in each cell type. Correlation coefficients for each pair were 0.96 in CD4+ T-cells, 0.97 in CD8+ T-cells, and 0.96 in CD14+ monocytes (**Figure 2a**). Distribution of CpG methylation in each cell type derived by TCA and MC-seq methods were almost identical. Compared to directly measured methylation, a small proportion of hypermethylated CpG sites with β>0.95 trended slightly higher using the TCA method (**Figure 2b**). These results suggest that TCA is a robust and effective deconvolution method. Accordingly, EWAS for HIV infection using TCA-deconvolution was performed.

**Figure 2.**
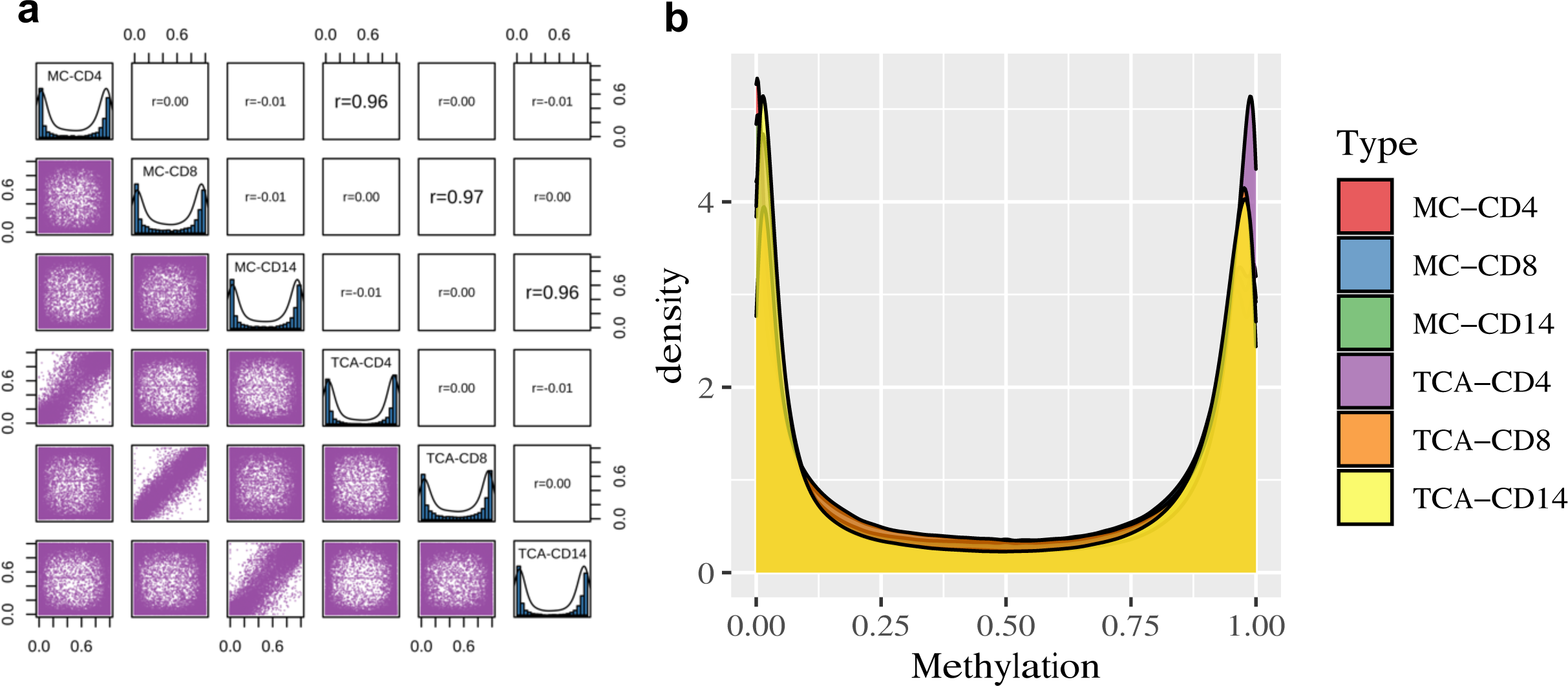
Benchmarking TCA-deconvoluted cell-type specific DNA methylation. (a) Comparison of methylation β-values for the top 10,000 most variable CpG sites between the deconvoluted and the directly measured methylation. β-values for each cell type were compared between three cell types [CD4+ T cells, CD8+ T cells, and monocytes (CD14+)]; (b) Distribution of genome-wide DNA methylome by the TCA-deconvoluted and MC-seq methods. MC: methylation capture sequencing; TCA: Tensor Composition Analysis.

### Cell type-based EWAS identified differentially methylated positions (DMPs) for HIV infection in men with HIV: the Veteran Aging Cohort Study

In the VACS cohort, the EWAS of HIV-infection using whole blood was carried out by applying a two-step regression model adjusting for age, self-reported race, cigarette smoking, alcohol use, ART adherence, HIV viral load, and the top 30 principal components (PCs) of DNA methylation. We identified 496 epigenome-wide significant (EWS) DMPs associated with HIV infection in the bulk tissue blood DNA methylome (false discovery rate, FDR<0.05) (**Figure 3a, Supplemental Figure 1a, sTable 2**). The significant DMPs included those previously reported by us and other groups. Examples include two previously replicated associations on *NLRC5* (cg16411857, t=-9.42, FDR=3.58E-14 and cg07839457, t=-9.05, FDR=5.68E-10) [17, 18 36]. Hypomethylation of *HCP5* that was previously linked to HIV infection also reached EWS in this study (cg18808777, t=5.61, FDR=7.81E-04) [18]. While replication of these well-validated CpG sites and genes is important, whether these DMPs originate from specific cell types or sub-groups of cell types is unknown.

**Figure 3.**
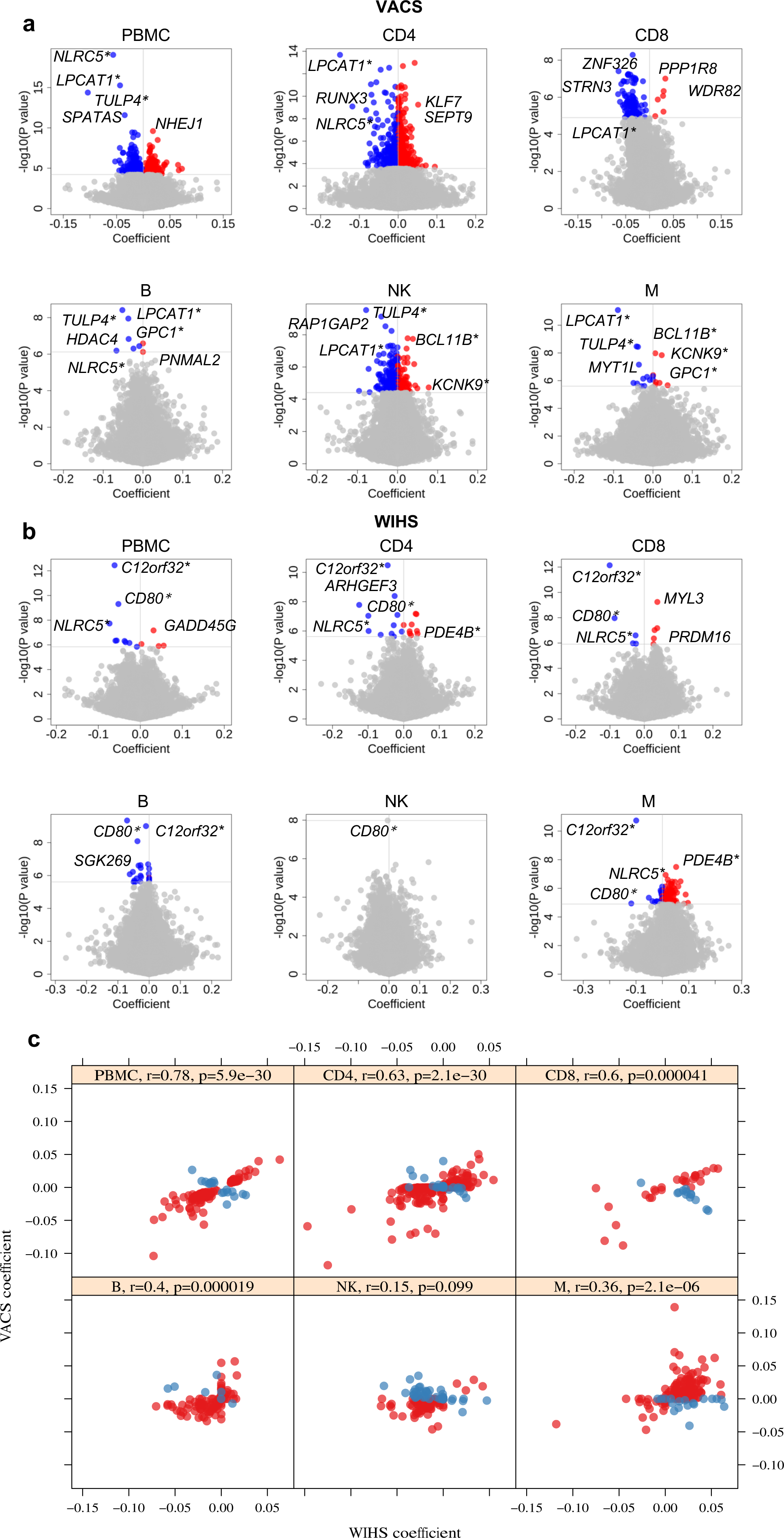
Summary of cell-type level EWAS in the VACS and WIHS cohorts in five cell types (CD4+ T, CD8+ T, B, Natural Killer, Monocyte). (a) Volcano plots for the VACS cohorts with top common and unique hyper- and hypomethylated gene-associated sites annotated, where PBMCs derive from whole blood samples. (b) Volcano plots for the WIHS cohort with similar annotations. (c) Correlation of significant DMPs between VACS and WIHS cohorts among the shared DMPs between the two cohorts. PBMC: peripheral blood mononuclear cell; EWAS: Epigenome-wide Association Study; VACS: Veteran Aging Cohort Study; WIHS: Women’s Interagency HIV Study; DMPs: Differential Methylation Positions. *Significant genes shared between at least two cell types.

At the cell-type level, we identified considerably more EWS DMPs across the five cell types than in whole blood: 2,208 in CD4+ T-cells, 106 in CD8+ T-cells, 8 in B cells, 317 in NK cells, and 21 in monocytes (**Figure 3a, Supplemental Figure 1b, Supplemental sTable 3**-**7**). The majority of DMPs in each cell type differed among the five cell types; a small number of DMPs were common in more than one cell type. For example, one DMP, *LPCAT1* cg16272981, was significant in four cell types (CD4+ T-cells: FDR=1.15E-07; B cells: FDR=0.0016; NK cells: FDR=0.005 and monocytes: FDR=3.29E-06). In total, 765 DMPs share more than one cell type in the same direction.

In CD4+ T-cells, differentially methylated CpGs were located in 1,407 genes. We found more hypomethylated (N=1,336) than hypermethylated (N=827) CpG sites in samples from PWH relative to uninfected controls. The HIV-associated loci previously reported in CD4+ T-cells were replicated in this cohort. *LPCAT1* cg16272981 showed the largest effect (16.1% less methylated in samples from PWH relative to PWoH). The top 30 DMPs were located in 15 genes (i.e. *LPCAT1, SLC17A9, RUNX3, KLF7, SEPT9, PEX14, NLRC5, SPOCK2, SPATAS, MYT1L, CAPN11, SEMA3G, BCL9, XYLT1*) (FDR=8.49-09∼ 5.21E-06). Some genes harbored multiple DMPs. For example, 4 EWS DMPs were located in *RUNX3*, a well-recognized tumor suppressor of gastric, colon and many other forms of solid tumors [37]. The majority of HIV-associated DMPs in CD8+ T-cells were hypomethylated. Six out of 8 DMPs in B cells were hypomethylated. *LPCAT1* cg16272981, also significantly enriched in CD4+ T-cells, showed the strongest EWS association in B cells (t=-5.97, FDR=0.003). Other significant DMPs were located on *TULP4, ETS1, KCNK9, STAT3, HDAC4*, and *GPC1*. Of note, 6 out of 8 DMPs were common between CD4+ T-cells and B cells except for *ETS1* and *GPC1*. In NK cells, 159 DMP sites overlapped with other cell types. *TULP4* cg02571055 was hypomethylated in B cells and the strongest EWS association in NK cells (t=-6.40, FDR=0.0001). Among 21 significant CpG sites in monocytes, the most significant DMP was *LPCAT1* cg16272981 (t=-6.97, FDR=3.29E-06), followed by *TULP4* cg02571055 (t=-5.99, FDR= 4.94E-04), which was also identified in B cells and NK cells. Thirteen DMPs were only observed in monocytes, including DMPs located in in *NOTCH4* and *IGSF9*.

### Cell-type EWAS for HIV infection in women with HIV: the Women’s Interagency HIV Study

The EWAS of women with HIV using PBMCs was carried out by applying the same regression model and adjusting for the same covariates as for the EWAS in the VACS. We identified 13 EWS DMPs associated with HIV infection (**Figure 3b, Supplemental Figure 1c, Supplemental sTable 8**). Consistent with the VACS sample, *NLRC5* cg07839457 was one of the most significant CpG sites (t=-5.76, p=1.85E-08). *NLRC5* cg16411857 showed near epigenome-wide significance (t=-4.51, p=8.95=-06). Other EWS DMPs were located in *C12orf32*, *CD80*, *GADD45G*, *TXNIP*, *TMEM49*, *SGK269*, *DUSP16*, *RAC2*, *TNIP3*, and *GLB1L2*.

For the cell-type level EWAS, we identified 153 significant DMPs among the 5 cell types: 20 for CD4+ T-cells, 10 for CD8+ T-cells, 22 for B cells, 1 for NK cells, and 100 for monocytes (all FDR<0.05) (**Figure 3b, Supplemental Figure 1d, Supplemental sTable 9**-**13**). Several DMPs are worthy of mention. *C12orf32* cg12051710 displayed the strongest association in four out of five cell types: CD4+ T-cells (t=-6.85; FDR=1.35E-05), CD8+ T-cells (t=-7.45, FDR= 2.95E-07), B cells (t= −6.28, FDR=2.02E-04), and monocytes (t=-6.94 p= 7.43E-06). *CD80* cg13458803 was hypomethylated in multiple cell types: CD8+ T-cells (t=-5.86, FDR=0.0001), B cells (t=-6.41, FDR=0.0002), and NK cells (t=-5.86, FDR=0.004).Of note, *NLRC5* cg07839457 was one of the top ranked DMPs in CD4+ T-cells in the WIHS (t=-5.78, FDR= 0.00537), consistent with the EWAS in the VACS.

### Concordance of DMPs for chronic HIV infection in the VACS and WIHS cohorts

The distinct demographic and clinical characteristics of the VACS and the WIHS may undermine whether the HIV-associated DMPs identified in these two cohorts are comparable and if the findings can be generalized to other studies. We conducted a correlation analysis of effect sizes for each DMP across data from bulk samples (whole blood, PBMC) and each cell type between the two cohorts (p<0.001). We found that effect sizes of the same CpG site between the two cohorts were highly correlated in bulk cells (r=0.784, p=5.86E-30) and in four out of five cell types (**Figure 3c**). At the cell type level, the correlation coefficient of DMPs from the VACS and WIHS were strongest in CD4+ T-cells (r=0.635, p=2.14E-30), followed by CD8+ T-cells (r=0.601, p=4.13E-05), B cells (r=0.396, p=1.89E-05), and monocytes (r=0.365, p=2.12E-06). The correlation of effect size in NK cells between the two cohorts was suggestive, but not significant (r=0.151, p=0.099). The directions of the correlations for the majority of DMPs (85.1%) were concordant between the two cohorts. The correlation analysis results show that DMPs for HIV infection in bulk and in individual cell types were largely consistent between the two cohorts; the results also underscored the potential value of EWAS meta-analysis of the two cohorts.

### EWAS meta-analysis by cell type identified common and specific DMPs for chronic HIV infection

Cell-type based EWAS meta-analysis (meta-EWAS) identified EWS DMPs in each of five cell types. Prior to performing the meta-EWAS, DMPs with significant heterogeneity (heterogeneity p<0.05) between the two cohorts were removed.

Meta-EWAS in bulk cells revealed 453 DMPs including top significant genes *NLRC5*, *LPCAT1*, *HCP5*, and *PSMB8* (**Figure 4a, Supplemental sTable 14**). Meta-EWAS by cell type identified 1,472 epigenome-wide DMPs in CD4+ T-cells, no DMPs in CD8+ T-cells, 159 DMPs in B cells, 198 DMPs in NK cells, and 422 DMPs in monocytes (**Figure 4b and 4c, Supplemental sTable 15-18**). In CD8+ T-cells, the DMPs uniquely identified in the VACS cohort showed opposite direction in the WIHS cohort, with the exception of the DMP in *C12orf32* and *MYL3*. Because the CpG sites with opposite effect were removed from the meta-analysis and no significant DMPs in CD8+ T-cells were identified due to high heterogeneity. Several genomic regions harbored DMPs that were common to more than one cell type (**Figure 5a**). Multiple loci on chromosome 3 (*ARHGEF3, DNAJB8, CCRL1, AHSG, CD80*), 5 (*LPCAT1*), chromosome 11 (*SHANK2*), chromosome 10 (*RUNX2*), chromosome 16 (*NLRC5*), and chromosome 20 (*SLC17A9*) were common in more than one cell type. Several top ranked CpG sites were located in genes encoding for transcription factors. For example, *ZNF326* (t=-5.92, FDR=0.002), *ZNF714* (t=-4.83; FDR=0.016), *ZNF90* (t=-4.45, FDR=0.042), and *ZNF76* (t=-4.44, FDR=0.044). Other DMPs were located in genes relevant to cancer such as *STRN3* cg18451035 (t=-5.56, FDR=0.004) and genes involved in innate immunity and metabolism such as *ENPP4* cg25606773 (t=-5.48, FDR=0.004) [38, 39].

**Figure 4.**
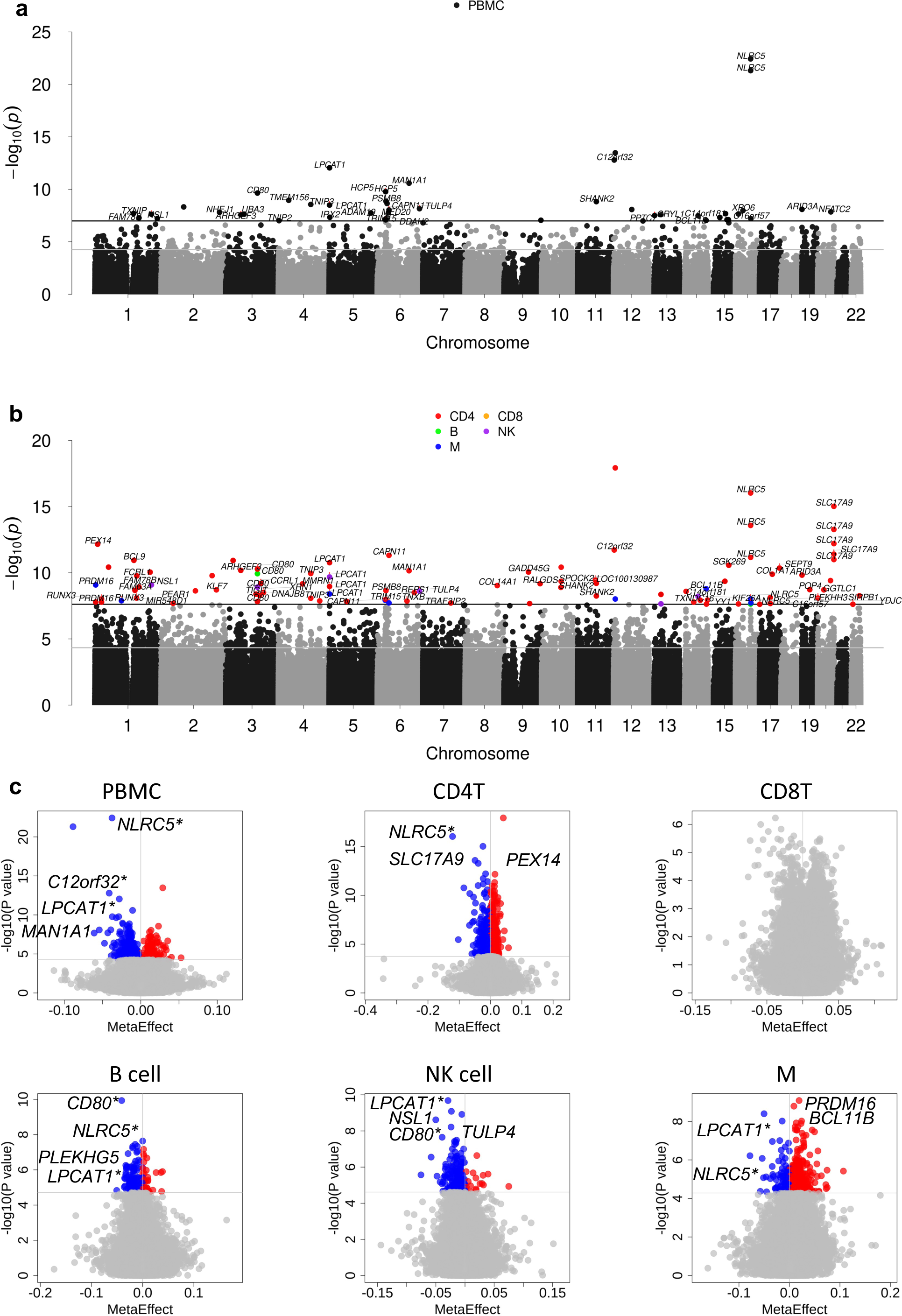
Summary of cell type level epigenome-wide meta-analysis of the combined Veteran Aging Cohort Study (VACS) and Women’s Interagency HIV study (WIHS) data. (a) Manhattan plot of epigenome-wide significant CpG sites prior to computational deconvolution of data into cell-type-specific methylation. (b) Manhattan plot of epigenome-wide significant CpG sites after computational deconvolution into cell-type-specific signals. (c) Volcano plots of hyper- and hypomethylated DMPs for HIV infection in each cell type following Meta-EWAS (Epigenome-wide Association Study). DMP: Differential Methylation Position. * Significant genes shared between at least two cell types.

**Figure 5.**
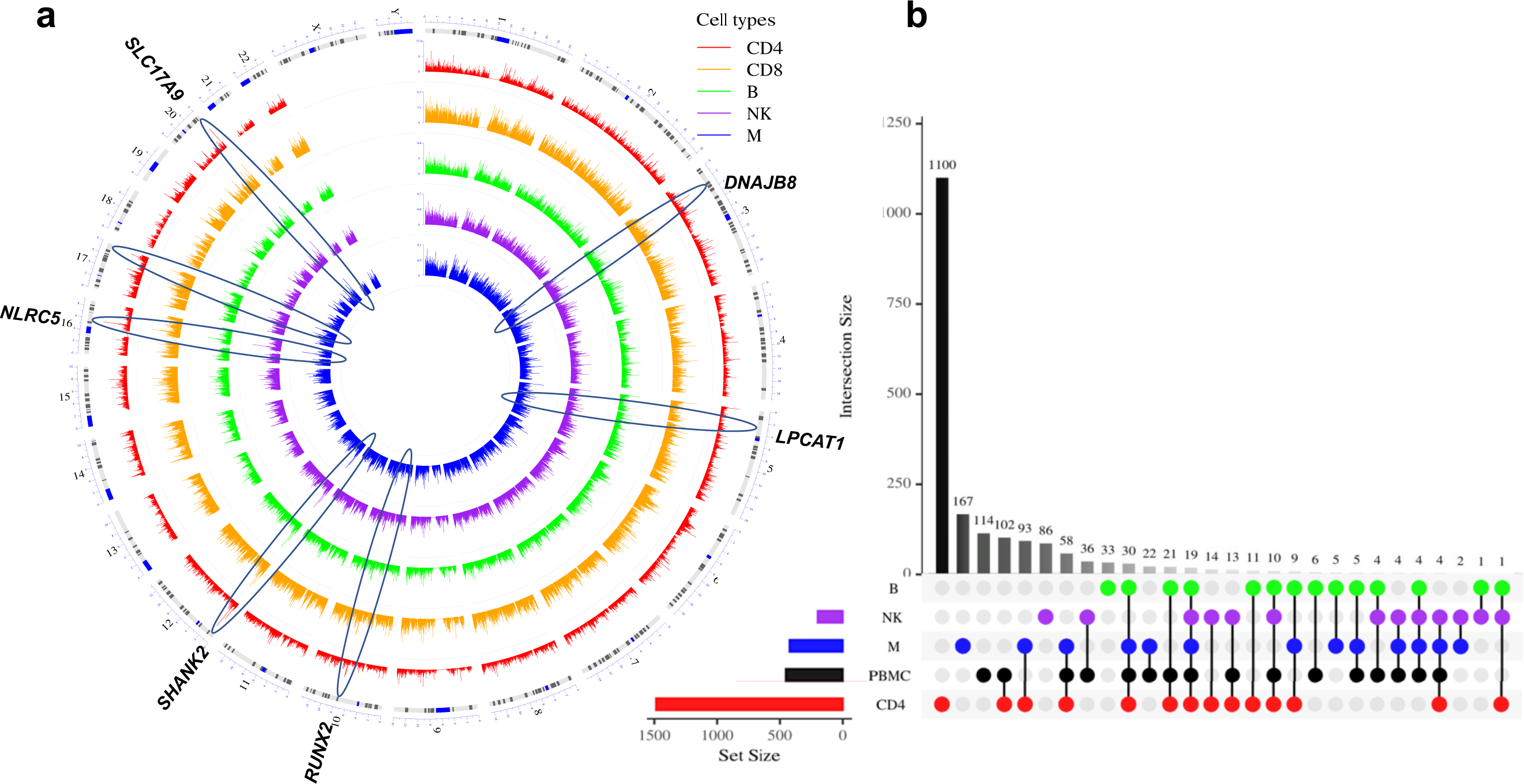
Common and distinct DMP profiles among cell types following cell type level meta-EWAS. (a) Stacked Manhattan plots displaying −log10(p) for CpG sites of each cell type. (b) Unique and common DMPs among five cell types and peripheral blood mononuclear cells. Overlap size represents the number of shared DMPs between the designated cell types. DMP: Differential Methylation Position.

Overall, the majority of DMPs (67%) from the meta-EWAS were unique to each cell type (**Figure 5b**). Cell-type specific DMPs accounted for 67.6% of DMPs in CD4+ T-cells, 20.8% in B cells, 42.4% in NK cells, and 39.3% in monocytes. Among lymphocytes, only 1.4% of DMPs overlapped between CD4+ T-cells and B cells, and 0.7% of DMPs between CD4+ T-cells and NK cells. The small number of overlapping DMPs may be because fewer DMPs were identified in B cells and NK cells. The results suggest that meta-EWAS revealed distinct DNA methylation modifications for HIV infection between cell types. Annotation of significant CpGs showed that the majority (40-76%) of CpG sites were located in gene bodies in each cell type. The proportion of DMPs located in promoter regions was greater in CD4+ T-cells (10%) than in other cell types, followed by monocytes (8%) (**Figure 6a**). The proportion of DMPs in CpG islands was also greater in CD4+ T-cells (**Figure 6b**).

**Figure 6.**
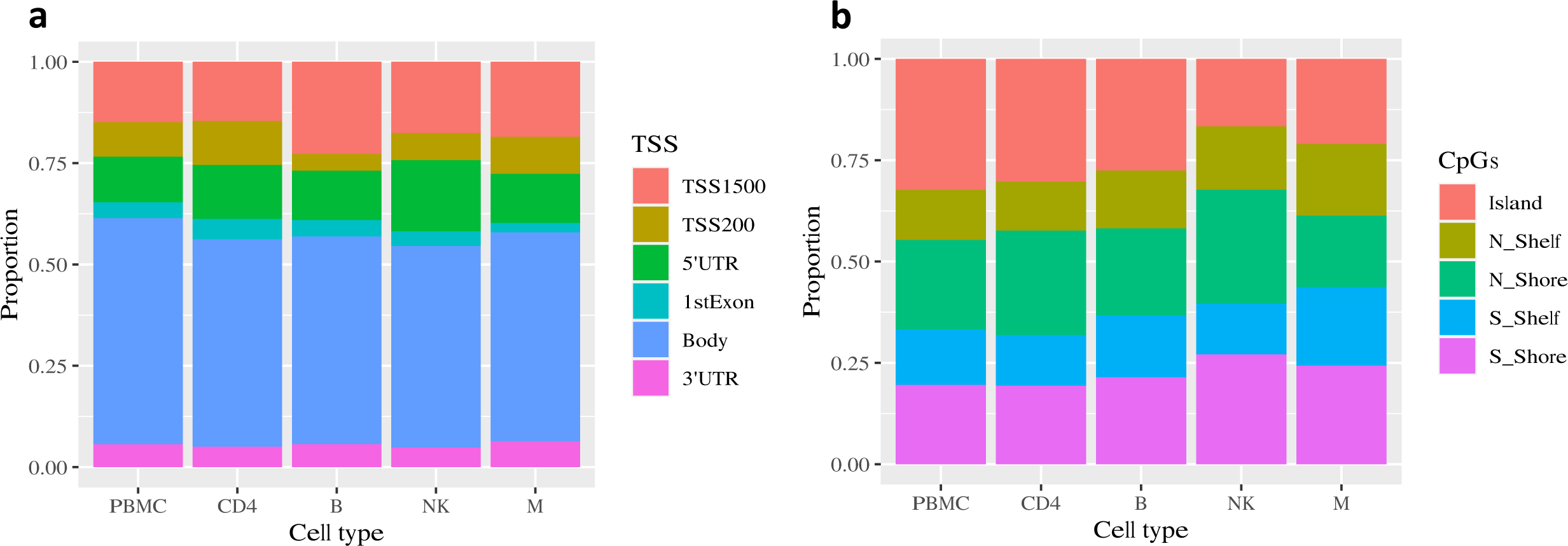
Characterization of epigenome-wide significant DMP from cell-type level meta-epigenome-wide association analysis. DMP: Differential Methylation Position.

### Cell-type Specific DMPs from meta-EWAS are overrepresented in hallmark genes for cancer

Previous studies have demonstrated that pathogen-induced epigenetic alterations cumulatively contribute to cancer development [40]. Using hallmark genes from the COSMIC Cancer Gene Census (databasehttps://cancer.sanger.ac.uk/census), we found significant overrepresentation of differential methylation of hallmark genes for cancer in CD4+ T-cells (FDR=1.01E-05), in B cells (FDR=0.008), in NK cells (FDR=0.003), and in monocytes (FDR=0.02). Several hallmark genes were EWS for HIV infection from the meta-EWAS in each cell type except CD8+ T-cells (**Table 1**).

**Table 1.**
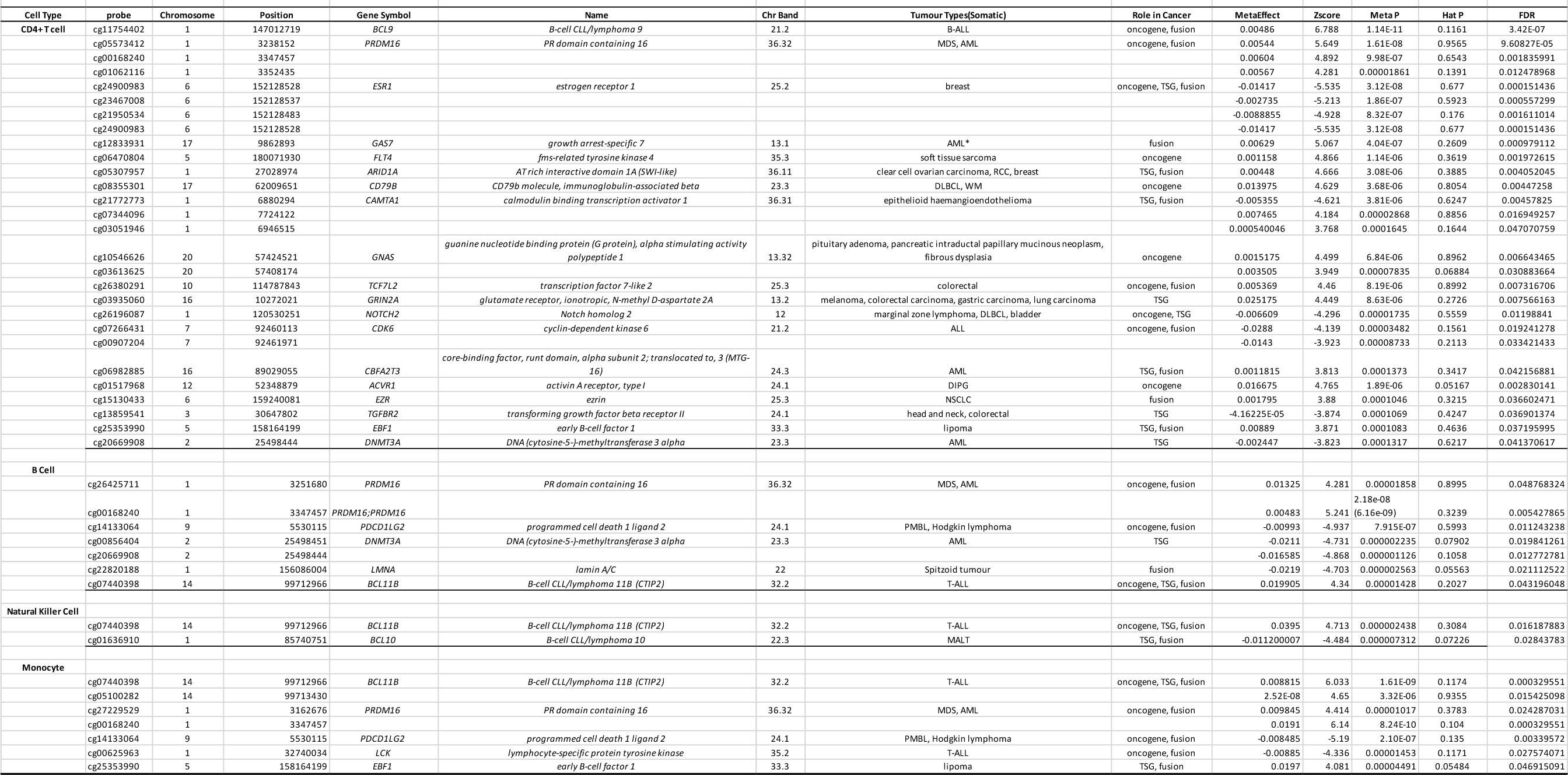
Overlap between hallmark genes for cancer and meta-EWAS identified significant genes for HIV infection

In CD4+ T-cells, 41 hallmark genes were differentially methylated for HIV infection (e.g. *BCL9* for B-ALL [41]*, GAS7* and *PRDM16 for* AML [42, 43]*, ESR1* for breast cancer [44]*, and GRIN2A* for colorectal, lung, and gastric carcinoma [45]) (**Figure 7**). Five hallmark genes were differentially methylated for HIV in B cells (*PRDM16* and *DNMT3A* for AML*, PDCD1LG2* for Hodgkin’s lymphoma*, LMNA* for spritzed tumor, and *BCL11B* for T-ALL). Three hallmark genes were differentially methylated for HIV in NK cells (*BCL11B* for T-ALL*, BCL10* for MALT*, and MAP3K7IP* for prostate cancer), and six in monocytes (*LCK* and *BCL11B* for T-ALL*, PDCD1LGD* for Hodgkin’s lymphoma*, PRDM16* for AML*, CACNA1D* for prostate cancer, and *EBF1* for B-ALL).

**Figure 7.**
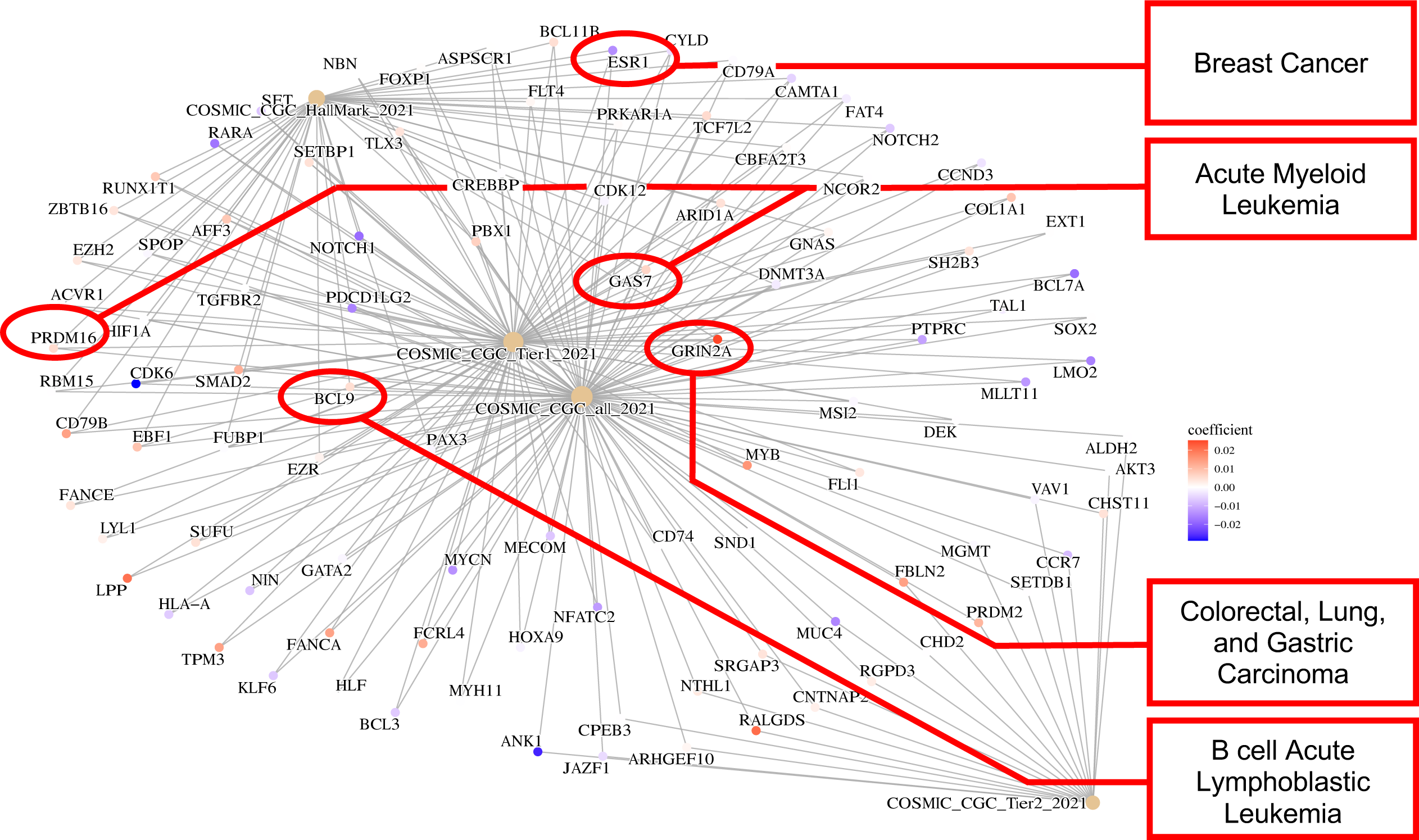
Gene enrichment analysis of significant DMPs for HIV infection in CD4+ T cells with Catalogue Of Somatic Mutations in Cancer (COSMIC) Gene Census Tier 1 database. CGC Tier 1 denotes the gene has cancer gene mutation patterns and evidence of functional impact reported in the literature. DMP: Differential Methylation Position.

In addition to the enrichment of hallmark genes for cancer, many of the HIV-associated DMPs in individual cell types are also reportedly involved in cancer. Some genes related to cancer harbored multiple CpG sites that all were hypomethylated in samples from PWH compared to PWoH Six DMPs on *SLC17A9* (cg14686919, cg01817521, cg00199007, cg04478428, cg26329715, cg00727912) were hypo-methylated in CD4+ T-cells, B cells, NK cells, and monocytes (**Table 1**). Notably, *SLC17A9* expression is associated with colorectal cancer [46]. We found 4 DMP sites on *RUNX3* that were hypomethylated (cg11585280, cg07236781, cg15498134, cg00147638). *RUNX3* is a tumor suppressor frequently deleted or transcriptionally silenced in cancers that encodes for one of the RUNX family proteins that are critical transcriptional regulators combining with transcription factor CBF-β in CD4+ T-cells [47]. Two *HCP5* DMPs were hypomethylated (cg18808777, cg25843003); *HCP5* is an oncogene associated with multiple cancers. DMPs in several genes from the *BCL* family were found in CD4+ T-cells, B cells, NK cells, and monocytes [48]. We found 5 EWS DMP sites on KLF7 that were hypermethylated. KLF7 affects cell proliferation and has been implicated in ovarian cancer progression [49]. Differential methylation of *KLF7* for HIV was previously linked to pancreatic ductal adenocarcinoma [5]. HIV-1 induced DNA methylation changes to CpGs in these genes, which have been reported in the literature to play a role in cancer development, may explain the increased prevalence of cancer in PWH.

## HIV-associated DMPs are enriched in gene sets for immunity and cancer biology

To better interpret the biological significance of HIV-associated DMPs in the aggregate, we carried out a gene set enrichment analysis. Among the set of hallmark genes, we found 20 significant pathways in CD4+ T-cells, 8 in B cells, 3 for NK cells, and 3 for monocytes (q<0.05). Multiple pathways were identified that were involved in immune evasion in multiple cancers (**Supplemental Table 19**). For example, the Kras pathway harbors a set of one of the most common oncogenic-driven mutations and genes that are targets of cancer therapeutics. In CD4+ T-cells, the Kras pathway contains HIV-associated hypomethylated DMPs in *IRF8, PSMBP8, PDCD1LG2, MYCN, and SNAP91* and hypermethylated DMPs in *BTBD3, CSF2*, and *NIN*. Interestingly, hypomethylation of *IRF8* was enriched in not only the Kras pathway, but also the allograft rejection pathway and the interferon γ response pathway. Multiple genes contained HIV associated DMPs in the allograft rejection pathway, including hypomethylated DMPs in *CD80, CD8A, CRTAM, HLA-DOA, PTPRC, STAT1, STAT4, and CD96* and hypermethylated DMPs in *MBL2* and *CD4*. HIV-associated DMPs in *STAT1, STAT4, HLA-A* and *TAP1* were common to both the allograft rejection pathway and the interferon γ response pathway (**Figure 8a**). The allograft rejection pathway and the Kras pathway are connected by HIV-associated DMPs located in the following hallmark genes for cancer: *HIF1A, CD79A, PDCD1LG2*, and *FLT4* (**Figure 8b**). The three above-mentioned pathways are involved in cancer development and progression.

**Figure 8.**
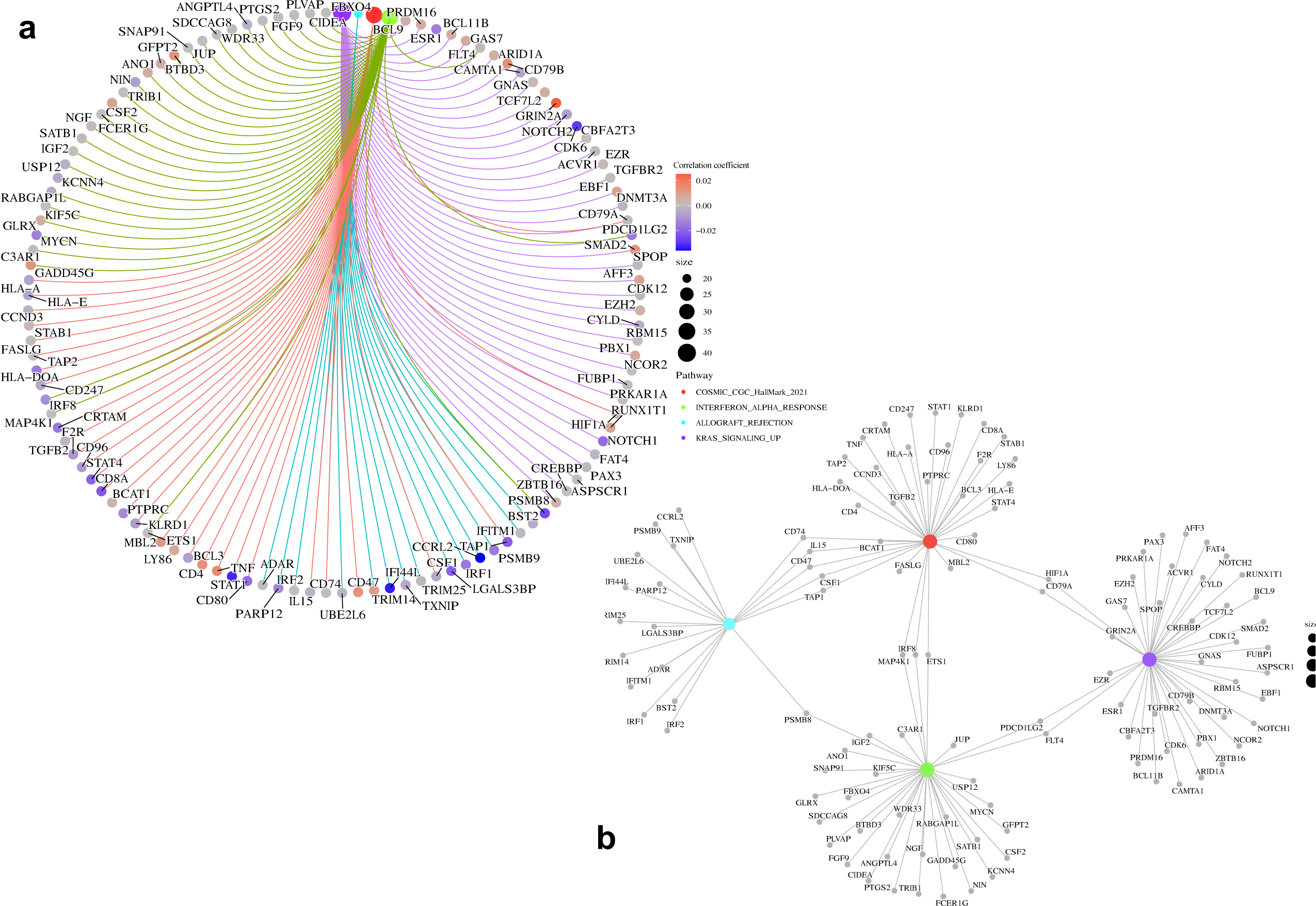
Gene enrichment analysis of DMPs in the genes enriched on hallmark gene pathways in CD4+ T cells. Notably, the allograft rejection, early estrogen response, interferon alpha response, interferon gamma response, and kras signaling pathways were significantly enriched. (a) Circos plot showing DMPs in four pathways; (b) Relationships of four significant pathways. DMP: Differential Methylation Position.

Gene set enrichment analysis for pathways in B cells mostly overlapped that in CD4+ T-cells. One significant pathway unique to B cells was the IL2-STAT5_SIGNALING pathway, in which 7 of 194 genes harbored HIV-associated DMPs. Gene set enrichment analysis for pathways in NK cells largely overlapped with those identified in B cells. One significant pathway unique in the NK cell was the HALLMARK_PI3K_AKT_MTOR_SIGNALING pathway, in which 7 of 104 genes harbored HIV-associated DMPs. Monocytes had one significant pathway: HALLMARK_PANCREAS_BETA_CELLS. Enrichment analysis from the KEGG database showed overall distinct patterns of significant biological pathways in each cell type **(Supplemental Figure 2)**. The results further underscore the striking enrichment of genes in cell type that feature DMPs for HIV-pathogenesis that are enriched for cancer development among PWH.

## Discussion

We identified DMPs for chronic HIV infection in five major immune cell types: CD4+ T-cells, CD8+ T-cells, B cells, NK cells, and monocytes. These include a number of previously reported DMPs associated with HIV infection. Despite differences in demographic and clinical characteristics between the two cohorts studied, we found a number of overlapping and highly concordant DMPs. The majority of DMPs identified in individual cell type meta-EWAS were unique to each cell type (67%). The occurrence of distinct profiles of HIV-associated DMPs among immune cell types highlights the importance of examining differences in DNA methylation profiles between individual cell types. Among the five cell types, the number of DMPs identified in CD4+ T-cells were three-to-ten-fold greater than the other four cell types, suggesting that epigenetic alteration in CD4+ T-cells plays a major role in chronic HIV-infection. More importantly, we found that genes that harbored HIV-associated DMPs are overrepresented in cancer biology. The identified genes were enriched among hallmark pathways of HIV pathogenesis and cancer. The results provide new insights into the epigenetic mechanisms of HIV that may underlie the increased risk for cancer in PWH.

As expected, we identified the largest number of HIV-associated CpG sites in CD4+ T-cells. Several previously reported genes involved in chronic HIV infection from CD4+ T-cells and from PBMC samples were replicated in this study. Of note, we observed differentially methylated CpG sites harbored in the genes involved in the Th1 signaling process (e.g. *RUNX3, STAT4*) in CD4+ T-cells. Several *TNF* CpG sites were reported to be hypermethylated in samples from PWH [50]. These CpG sites were also hypermethylated in the present study. A noteworthy hypomethylated DMP identified in the present study is *PEX14* cg25310676. PEX14 is involved in the control of oxidative stress and is targeted by HIV Env-mediated autophagy [51]. Expression of *PEX14* was decreased in HIV-infected CD4+ T-cells and contributed to CD4+ T-cell apoptosis [7]. Compared to CD4+ T-cells, a much smaller number of DMPs were identified in the other four cell types, including monocytes. This observation is in line with previous reports showing that different numbers of DMPs between acute and chronic HIV infection are observed in CD4+ T-cells and monocytes. During acute HIV infection, the number of DMPs are 10-fold greater in monocytes than in CD4+ T-cells [16]. In SIV-infected macaques and African green monkeys, only 0.5% of DMPs overlapped in CD4+ T-cells between acute and chronic HIV infection stages. This evidence suggests that distinct profiles of DNA methylation modification may occur in the different stages of HIV infection and in different cell types.

The fact that many HIV-associated DMPs and associated genes are also involved in cancer among the four cell types is intriguing. One possibility is that HIV-1 directly induces maladaptive changes in epigenetic regulation of oncogenes. For example, several BCL family genes were significantly associated with HIV infection, *BCL9* in CD4+ T-cells, and both *BCL11B and BCL2L2* in CD4+ T-cells, B cells, NK cells, and monocytes. The BCL family plays a crucial role in the development, proliferation, differentiation, and subsequent survival of T cells and is associated with multiple cancers. *BCL9* functions in cell-cell communication in colorectal cancer [52]. *BCL11B* encodes for a protein that is a transcriptional repressor and is regulated by the NURD nucleosome remodeling and histone deacetylase complex. *BCL11B* is a hallmark of B-cell CLL/Lymphoma. *BLC2L2* acts as an apoptotic regulator and is linked to multiple cancers including liver cancer, lung cancer, and breast cancer [53]. On the other hand, evidence shows that chronic inflammation is involved in pathogen-induced DNA methylation changes resulting in an “epigenetic field defect” for oncogenesis. For example, our results show that several proinflammatory genes (e.g. *TNF, IGFBPL1*) were differentially methylated in PWH compared to PWoH. Increased inflammation is a hallmark of chronic HIV infection. Whether chronic HIV-1 results in DNA methylation of inflammatory genes contributing to cancer warrants further study. The overrepresentation of HIV-1 integration in cancer genes has been reported previously [54]. In PWH on suppressive ART, a large proportion of persisting proviruses are found in proliferating cells. One possible mechanism is to promote the proliferation and survival of latently HIV-infected cells, which in turn benefits HIV-1 persistence and reservoir expansion thereby frustrating attempts to eradicate the virus. Such interactions between HIV-1 and the host epigenome may point to underlying mechanisms of cancer development in PWH.

Overall, we found highly concordant effect sizes among the shared DMPs in four of five cell types between two distinct cohorts. The observation highlights the common biological pathways in ancestrally heterogenous populations among men and women with HIV. On the other hand, we observed a subset of DMPs that differed between the two cohorts. We speculate that the DMPs identified that differed between the two cohorts are likely due to different statical power and different rates of HIV viral suppression between the two cohorts. The sample size of the VACS cohort is over twice that of the WIHS cohort. More importantly, participants in the VACS cohort had higher HIV viral loads than participants in the WIHS cohort. Although HIV viral load was adjusted for in each EWAS, residual effects on DNA methylation cannot be ruled out. Another possibility is biological differences in HIV infection between men and women. Sex differences in HIV infection are observed in clinical settings. Women appear better able to control HIV-1 replication compared to men, typically having lower HIV viral loads, and higher CD8+ and CD4+ T-cell counts [55]. However, the rate of progression to AIDS between men and women are similar, suggesting that immune and inflammatory activation are higher among women [55]. The underlying reasons for the observed differences in CD8+ T-cell epigenome are unclear. One study showed that an increased CD8+ T-cell count may be due to an enhanced capacity to respond to the IL12 cytokine in women compared to men, which leads to more effector cell differentiation [56]. Treatment naïve women with HIV-1 had significantly higher CD8+ T-cell activation than men, which appears mediated by interferon α in response to Toll-like receptor 7 [57]. Future studies including both sexes and addressing confounding factors are warranted to investigate potential sex differences in the host genome among PWH.

We acknowledge several limitations to this study. Cell type proportion was estimated based on DNA methylation, not cell count, which could result in inaccurate deconvolution of DNA methylation for individual cell types. While the results suggest the effect would be modest, TCA-deconvoluted DNA methylation profiles in each cell type may differ between the two cohorts due to differences in biospecimen collection. For the VACS cohort, cell-type DNAm was deconvoluted from whole blood that included granulocytes while cell-type DNAm from the WIHS cohort was deconvoluted from PBMCs, which excludes granulocytes. Computationally identified significant DMPs warrant confirmation in sorted cell types. Finally, only a small proportion of CpG sites in the methylome were investigated in this study (i.e., 450K and EPIC commercial arrays). Future studies to expand the number of CpG sites using a sequencing platform to comprehensively profile the methylome for chronic HIV infection are warranted.

In summary, leveraging a computational deconvolution approach, we identified cell-type level DPMs associated with HIV infection. The findings were enriched for genes involved in both HIV pathogenesis and cancer pathology, which underscore the important mechanisms of HIV persistence and comorbid cancers. The significant genes may be therapeutic targets for HIV disease and other comorbid medical diseases.

## Methods

### Sample characteristics and DNA methylation profiling

The VACS is a nationwide longitudinal veteran cohort including PWH and PWoH to study HIV infection and disease progression. A total of 702 samples from the VACS Biomarker Cohort, a subset of the entire VACS, were included in the analysis (**Supplemental Material**). The majority (86%) of the VACS sample were of African (African American/Black; AA) ancestry and all samples were collected from male participants. Clinical data and specimens used in this manuscript were collected by the Women’s Interagency HIV Study (WIHS), now the Multicenter AIDS Cohort Study (MACS)/WIHS Combined Cohort Study (MWCCS) [58] **(Supplemental Material)**. WIHS included 245 samples from PWH and 187 uninfected controls from diverse ancestral populations and all samples were collected from female participants. Demographic and clinical characteristics are presented in **Supplemental sTable 1**.

Methylation of bulk DNA samples extracted from whole blood in the VACS was profiled using Illumina HumanMethylation 450K Beadchip. Methylation of bulk DNA samples extracted from PBMCs in the WIHS was profiled using Illumina HumanMethylation EPIC Beadchip. A total of 408,366 CpG sites were common to both the 450K and EPIC arrays, which were used to deconvolute bulk DNA methylation data to five cell types by TCA. More information about DNA methylation quality control and deconvolution are presented below in the Methods and in **Supplementary Material**.

### Capture Methylation Sequencing

Three cell types, CD4+ T-cells, CD8+ T-cells, and monocytes, were isolated from 4 PBMC samples using a magnetic bead-based method [59]. DNA was extracted from each isolated cell type. Methylation sequencing target enrichment library preparation was performed per manufacturer protocol (Agilent). Samples were sequenced using 100bp paired-end sequencing on an Illumina HiSeq NovaSeq according to Illumina standard protocol. Detailed quality control and data processes are presented in **Supplementary Material**. CpG sites were annotated using Homer annotatePeaks.pl, including intergenic, 5’UTR, promoter, exon, intron, 3’UTR, transcription start site (TTS), and non-coding categories. CpG island, shore, shelf, and open sea annotation was defined by locally developed bash and R scripts based on genomic coordinates (hg19) of CpG islands from the UCSC genome browser. CpG shore was defined as up to 2 kb from CpG islands and CpG shelf was defined as up to 2 kb from a CpG shore. Methylation CpG sites on the X and Y chromosomes were removed for subsequential analyses.

### Deconvolution of DNA methylation from bulk cells to five cell types

To deconvolute bulk methylation of each CpG to specific cell types, TCA requires a DNA methylation data matrix in heterogeneous cells and cell type proportions for each sample in the cohort. We first estimated the proportion of six cell types from the methylation of whole blood in the VACS cohort (CD4+ T-cells, CD8+ T-cells, B cells, NK cells, monocytes, granulocytes) and from the PBMCs in the WIHS cohort (CD4+ T-cells, CD8+ T-cells, B cells, NK cells, monocytes). Because the approach to generating PBMC in the WIHS cohort results in the near-total depletion of granulocytes, we excluded granulocytes from the analyses to provide a consistent set of cell-type specific epigenome profiles shared between the two cohorts. The estimated proportions of each cell type in each cohort were similar except for CD4+ T-cells, for which the proportion in the VACS cohort was greater than CD4+ T-cells in the WIHS cohort (**Supplemental Figure 3**). We used TCA in the R environment to estimate methylation beta values at each CpG site for each cell type. A total of 408,366 CpG sites were deconvoluted to five cell types using either a whole blood methylation matrix (VACS) or from PBMC methylation matrix (WIHS). Our results showed that TCA deconvoluted methylation in individual cell types robustly removed cell type confronting effects in (**Supplemental Figure 4**).

### Comparison between TCA-deconvoluted and methylation capture sequencing-based methylation beta values in three cell types

We validated the accuracy of the TCA-derived estimates of DNA methylation at each CpG site by comparing the TCA-derived beta value with DNA methylation beta estimates measured directly using capture sequencing. We selected the top 10,000 most variable CpG sites among the samples for this comparison, which was performed using Pearson correlation analysis with significance set at p<0.05.

### Cell-type based epigenome-wide association analysis

We performed a cell-type based EWAS for HIV-infection using TCA-deconvoluted methylation beta values for each cell type. In each cell type, we conducted a two-step regression analysis using the strategy proposed by Lehne *et al [60]*. The first regression model addressed global covariates that may confound the association of methylation with HIV infection. We first estimated the residual β using regression model (<u>1</u>):

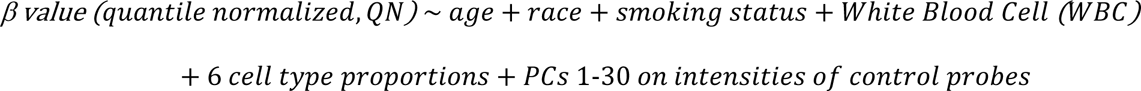

We then performed a second PCA on the resulting regression residual β values and regressed out the first 5 PCs to further control for unmeasured confounders in the regression model (2).

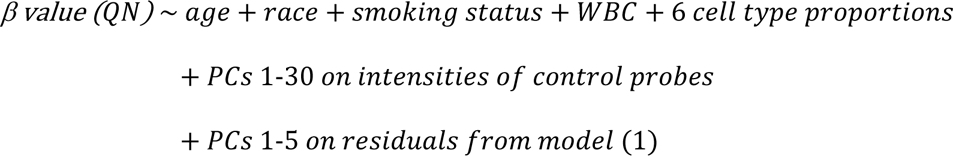

Significance was set at a false discovery rate (FDR)<0.05. To further confirm the correction of global confounders, we performed Pearson correlation analysis between the first 30 PCs on residual methylation from model (1) and batch, demographic, clinical, and 6 cell type confounders (**Supplemental Figure 4**). Cutoff of correlation analysis was set at p<0.05

### Correlation analysis of HIV-associated CpG sites between the two cohorts

In each cell type, we selected CpG sites with FDR<0.05 in the VACS and with a nominal p<0.05 in the WIHS for correlation analysis. The rationale for the significance cut off chosen for the WIHS cohort is to treat it as a replication cohort. Pearson correlation of the effect sizes at each resulting CpG between the two cohorts was performed. The significance threshold was set at p<0.05. We also compared the direction of effect of each CpG between two cohorts.

### Cell type-based Meta-EWAS

We conducted an EWAS meta-analysis for each cell type by combining the data from the VACS and WIHS samples. Effect sizes and p-values for each probe were obtained from analyses in the VACS and WIHS samples, respectively. We performed inverse-variance weighted meta-analysis, with scheme parameters of sample size and standard error as implemented in the METAL program, combining summary statistics from the two sample sets. We investigated heterogeneity between the two samples using the *I*^2^-statistic. CpG sites with *I*^2^>50% and heterogeneity p<0.05 were excluded from subsequent analysis.

### Enrichment of hallmark genes for cancer

We performed an enrichment analysis of HIV-associated genes among 736 cancer hallmark genes from the Cancer Gene Census database. CpG sites with FDR<0.1 from the cell type based meta-EWAS were mapped to the nearest gene. The set of genes that met the above criteria were used to test whether the gene set was significantly overrepresented among the hallmark genes for each cell type at FDR<0.05.

### Gene set enrichment analysis

Genes adjacent to CpG sites with FDR<0.1 in meta-EWAS for each cell type were selected for gene set enrichment analysis. We focused on the hallmark gene sets from the Molecular Signature Database (https://www.gsea-msigdb.org/gsea/msigdb/). Enrichment analysis using GO and KEGG annotations were also performed.

## Declarations

### Ethics approval and consent to participate

The study was approved by the committee of the Human Research Subject Protection at Yale University and the Institutional Research Board Committee of the Connecticut Veteran Healthcare System, and by the participating clinical sites. All subjects provided written consent. The views and opinions expressed in this manuscript are those of the authors and do not necessarily represent those of the Department of Veterans Affairs or the United States government.

### Availability of data and materials

Demographic and clinical variables and DNAm data for the VACS samples were submitted to GEO dataset (GSE117861) and are publicly available. All codes for analysis are also available upon a request to the corresponding author.

### Competing interests

VCM has received investigator-initiated research grants (to the institution) and consultation fees (both unrelated to the current work) from Eli Lilly, Bayer, Gilead Sciences, and ViiV. The remaining authors declare that they have no competing interests.

## Funding

The project was supported by U10 AA013566-completed from National Institute on Alcohol Abuse and Alcoholism (VACS award for which the majority of our cohort data is based) U24 AA020794 (COMpAAAS coordinating center), U01 AA020790, and R03DA039745, R01DA038632, R01DA047063, R01DA047820 from the National Institute on Drug Abuse.

COMpAAAS/Veterans Aging Cohort Study, a CHAART Cooperative Agreement, is supported by the National Institutes of Health: National Institute on Alcohol Abuse and Alcoholism (U24-AA020794, U01-AA020790, U01-AA020795, U01-AA020799; U10-AA013566-completed) and in kind by the US Department of Veterans Affairs. In addition to grant support from NIAAA, we gratefully acknowledge the scientific contributions of Dr. Kendall Bryant, our Scientific Collaborator. Additional grant support from the National Institute on Drug Abuse R01-DA035616 is acknowledged.

MWCCS (Principal Investigators): Atlanta CRS (Ighovwerha Ofotokun, Anandi Sheth, and Gina Wingood), U01-HL146241; Bronx CRS (Kathryn Anastos and Anjali Sharma), U01-HL146204; Brooklyn CRS (Deborah Gustafson and Tracey Wilson), U01-HL146202; Data Analysis and Coordination Center (Gypsyamber D’Souza, Stephen Gange and Elizabeth Golub), U01-HL146193; Chicago-Cook County CRS (Mardge Cohen and Audrey French), U01-HL146245; Northern California CRS (Bradley Aouizerat, Jennifer Price, and Phyllis Tien), U01-HL146242; Metropolitan Washington CRS (Seble Kassaye and Daniel Merenstein), U01-HL146205; Miami CRS (Maria Alcaide, Margaret Fischl, and Deborah Jones), U01-HL146203; UAB-MS CRS (Mirjam-Colette Kempf, Jodie Dionne-Odom, and Deborah Konkle-Parker), U01-HL146192; UNC CRS (Adaora Adimora), U01-HL146194. The MWCCS is funded primarily by the National Heart, Lung, and Blood Institute (NHLBI), with additional co-funding from the Eunice Kennedy Shriver National Institute Of Child Health & Human Development (NICHD), National Institute On Aging (NIA), National Institute Of Dental & Craniofacial Research (NIDCR), National Institute Of Allergy And Infectious Diseases (NIAID), National Institute Of Neurological Disorders And Stroke (NINDS), National Institute Of Mental Health (NIMH), National Institute On Drug Abuse (NIDA), National Institute Of Nursing Research (NINR), National Cancer Institute (NCI), National Institute on Alcohol Abuse and Alcoholism (NIAAA), National Institute on Deafness and Other Communication Disorders (NIDCD), National Institute of Diabetes and Digestive and Kidney Diseases (NIDDK), National Institute on Minority Health and Health Disparities (NIMHD), and in coordination and alignment with the research priorities of the National Institutes of Health, Office of AIDS Research (OAR). MWCCS data collection is also supported by UL1-TR000004 (UCSF CTSA), P30-AI-050409 (Atlanta CFAR), P30-AI-050410 (UNC CFAR), and P30-AI-027767 (UAB CFAR). VCM received support from the Emory CFAR (P30 AI050409).

## Authors’ contributions

XZ was responsible for data analysis and manuscript preparation. YH and CY were involved analytical strategy and supervised data analysis. RV contributed to manuscript preparation. ACJ provided DNA samples, clinical data, and contributed to interpretation results and manuscript preparation. MHC contributed to data collection. VM involved in interpretation results and manuscript preparation. BA and KX contributed equally to study design, interpretation findings, and manuscript preparation. All co-authors reviewed and approved the manuscript.

## Acknowledgements

The authors appreciate the support of the Veteran Aging Study Cohort Biomarker Core, Women’s Interagency HIV Study, and Yale Center of Genomic Analysis. The views and opinions expressed in this manuscript are those of the authors and do not necessarily represent those of the Department of Veterans Affairs or the United States government. This work uses data provided by patients and collected by the VA as part of their care and support.

