## Supplemental Figure for "Cell-type specific EWAS identifies genes involved in HIV pathogenesis and oncogenesis among people with HIV infection"

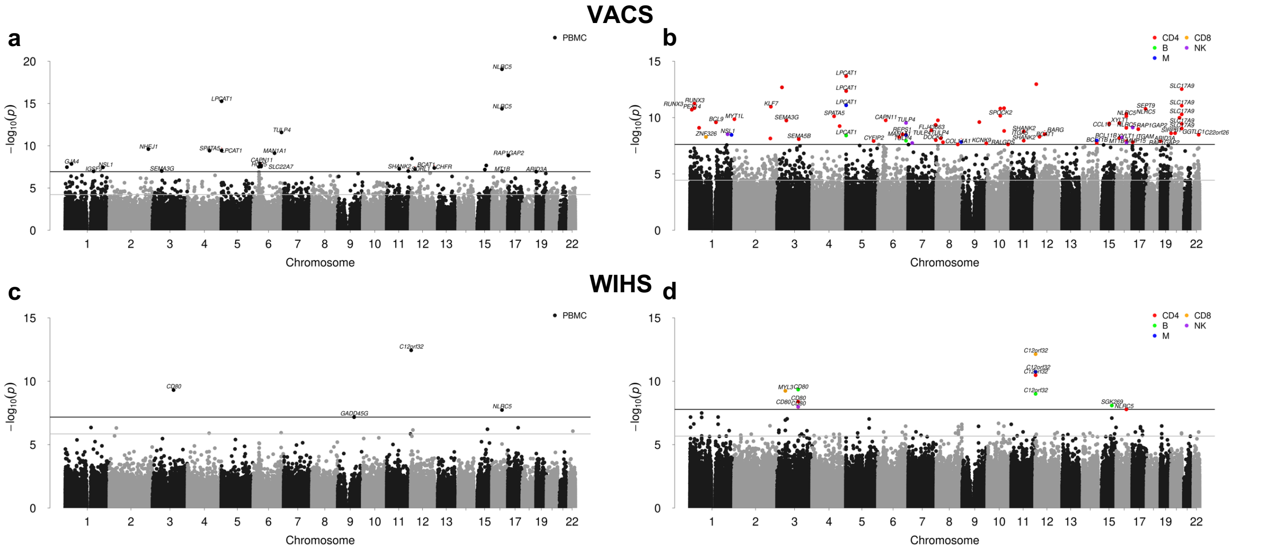


Supplemental Figure 1. Summary of EWAS prior to and following computational deconvolution in each cohort. (a) VACS, EWAS on HIV infection in whole blood. (b) VACS, EWAS on HIV infection in 5 individual cell types. (c) WIHS, EWAS on HIV infection in PBMCs. (d) WIHS, EWAS on HIV infection in 5 individual cell types. EWAS: Epigenome-wide Association Study; VACS: Veteran Aging Cohort Study; WIHS: Women's Interagency HIV Study.

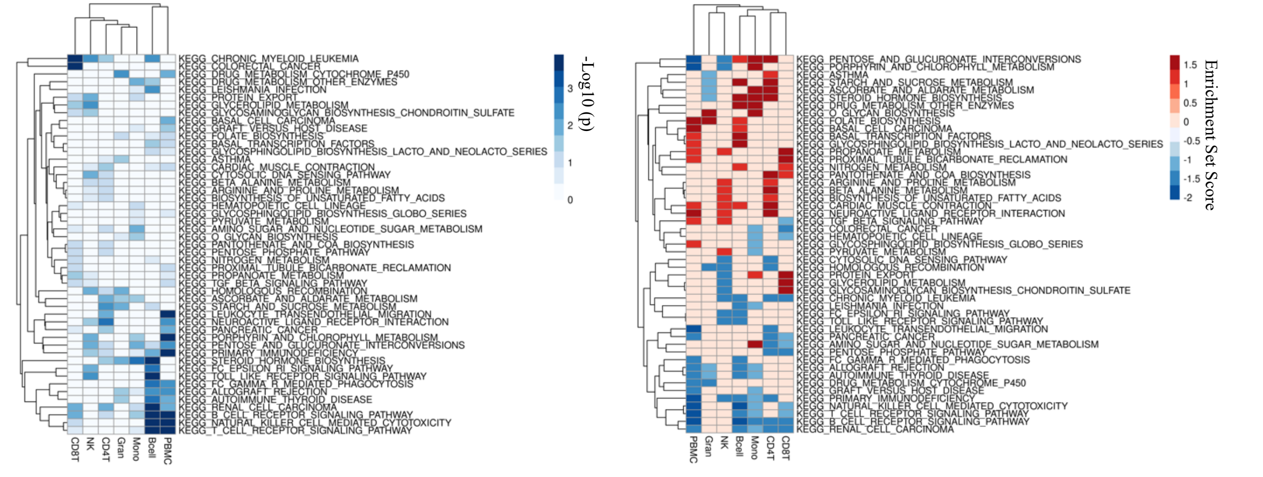


Supplemental Figure 2. KEGG (Kyoto Encyclopedia of Genes and Genomes) pathway analysis for each cell type.

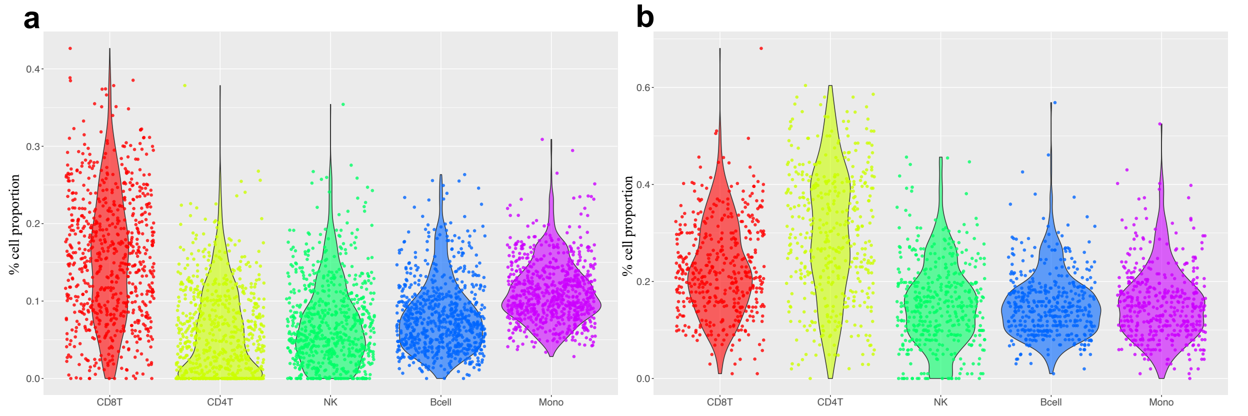


Supplemental Figure 3. Cell proportion estimation prior to TCA in both VACS and WIHS cohorts. (a) VACS cell proportion. (b) WIHS cell proportion. Granule cell type was removed in the sequential cell-type based EWAS analyses. EWAS: Epigenome-wide Association Study; VACS: Veteran Aging Cohort Study; WIHS: Women's Interagency HIV Study.

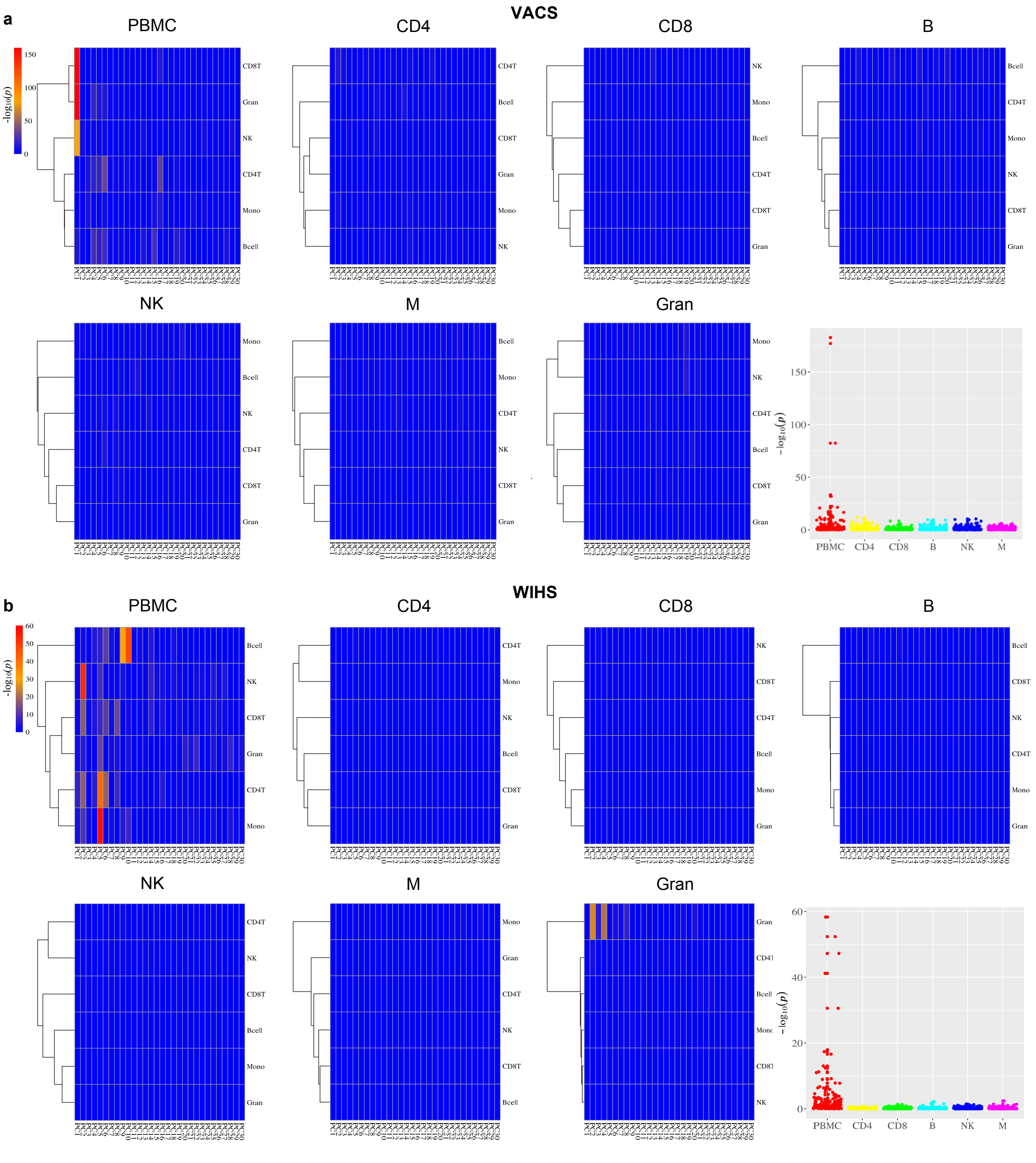


Supplemental Figure 4. Correlation analysis of top 30 Principle Components (PCs) on DNA methylation and cell type proportion in bulk cells and TCA-deconvoluted cell types. (a) VACS: -log$_{10}$(p) plots for whole blood methylation and for six individual cell type methylation. A jitterplot displays -log$_{10}$(p) value excluding granule cells. (b) WIHS: -log$_{10}$(p) plots for PBMC methylation and for six individual cell type methylation. A jitterplot displays -log$_{10}$(p) value for five cell types. PBMC: peripheral blood mononuclear cell; TCA: Tensor Composition Analysis; VACS: Veteran Aging Cohort Study; WIHS: Women's Interagency HIV Study.
