## Supplemental Material for "Cell-type specific EWAS identifies genes involved in HIV pathogenesis and oncogenesis among people with HIV infection"

Corresponding to

Ke Xu, MD, PhD

Bradley E Aouizerat, Ph.D.

### Supplemental Material

#### *Study cohorts*

*The Veteran Aging Cohort Study (VACS)*[1]: VACS is a nation-wide longitudinal cohort study in the United States to study medical and psychiatric comorbidities among people living with HIV (PWH). Since 1997, over 40,000 HIV-positive veterans and 80,000 age/race/site matched HIV-negative control participants are enrolled (https://www.vacsp.research.va.gov/CSPEC/Studies/INVESTD-R/Veteran-Aging-Cohort-Study.asp). A subset of the cohort participants provided blood specimens between 2005 and 2007 and consented for DNA analyses. At the time of sampling, the subjects were middle aged (>80% between 41 and 64 years of age). Most are men (95% male) and of African American ancestry (68%).

*The Women's Interagency HIV Study (WIHS)*[2]: WIHS is a prospective observational cohort study for the HIV epidemic, the natural history of HIV infection, and pathogenesis of HIV progression among women living with HIV. It was established in 1993. The WIHS is the largest and oldest ongoing prospective cohort study of women with and at risk for HIV infection in the world. The data collection included demographics, clinical variables, and biospecimens for HIV disease and comorbidities. Participants were African Americans (72%), European Americans was 11%, and 14% of Hispanic. The WIHS also recruited matched HIV-seronegative participants. Among Women with HIV, 97% were on antiretroviral therapy and 31% of them were not HIV-1 suppressed. Recently, WIHS was combined with the Multicenter AIDS Cohort Study (MACS) cohort, which is a longitudinal data and specimen-collection study for 35 years for men[3]. The combined cohort (MWCCS) included 2,115 women and 1,901 men with a median age of 56 years (interquartile range, 48–63); 62% are PWH[4].

***DNA methylation microarray data quality control (QC)***

DNA methylation was profiled using Illumina HumanMethylation 450K (for the VACS samples) or EPIC BeadChips (for the WIHS samples). The *minfi* R package (version 1.18.1) was used to retrieve both Illumina Infinium 450K and Illumina Infinium MethylationEPIC microarray raw data from the idat files. Normalization was performed for the HM450K and EPIC methylation data separately. For 450K dataset, quantile normalizations of 6 separated parts of intensity values were performed following the recommendations by Lehne et al[5]. For EPIC dataset, the ssNoob method was conducted[6]. The probe on Y chromosomes were used to evaluate the detection p value cutoff. P values < 1e−12 and 1e-8 were set as a detection p value threshold to improve the quantification of methylation intensities for 450K and EPIC, respectively. CpG sites on the sex chromosomes (X: N=11,232 CpGs, Y: N=416 CpGs) and the CpG sites with annotated single nucleotide polymorphism (SNP) within 10 bp (N=47,790) were excluded from the analysis. Previously identified cross-reactive probes were also excluded [7]. A total of 437,722 CpG sites in HM450K and 846,604 CpG sites in EPIC remained for the analysis. Three samples with a call rate <98% were also excluded. We compared the predicted sex with the self-reported sex and mis-matched samples were excluded.

#### ***DNA methylation capture sequencing (MC-seq) quality control***

DNA methylation was profiled by using Agilent SureSelectXT Methyl-seq Target Enrichment panels. Quality control was conducted following the standard procedure as previously described by Wreczycka et al. Quality of sequence raw data was examined by using FastQC (ver. 0.11.8). Adapter sequences and fragments at 5′ and 3′ with poor quality (phred score<20) were removed by Trim_galore (ver. 0.6.3_dev). We further applied Bismark pipelines[8] (ver. v0.22.1_dev) to align the reads to the bisulfite human genome (hg19) with default parameters. Quality- trimmed paired-end reads were transformed into a bisulfite converted forward strand version (CT conversion) or into a bisulfite-treated reverse strand (GA conversion of the forward strand). Duplicated reads were removed from the Bismark mapping output by *deduplicate_bismark* and CpG, CHG, and CHH (where H=A, T, or C) were extracted by *bismark_methylation_extractor*. NormalizeCoverage function in R package methyKit was performed. Beta values were calculated by Methy / (Methy + Unmethy + offset). The CpG data with coverage (also known as read depth) < 10 was labeled as NA. All the CpGs with call rate < 0.95 were excluded. Samples with call rate < 0.9 were also excluded. We used Homo software annotatePeaks.pl for gene annotation of CpG sites. A total of 1,841,043 CpG sites were used for the analysis.

#### ***Deconvoluting cell type methylation using Tensor Composition Analysis (TCA)***

Cell type methylation was deconvoluted for CD4+ T cells, CD8+ T cells, B cells, Natural Killer cells, Monocytes following the method by Rahmani et al[9]. The TCA method[10] is based on a hypothesis that the cell-type specific methylation levels of an individual are coming from a distribution that is shared across individuals in the population. Thus, the

TCA method is built to learn the unique cell-type specific methylomes of each individual sample from its bulk data. Briefly, a two-step pipeline was utilized for the estimation of differential DNAm in TCA. The first step involved a joint model that tested for evidence of differential DNAm at a CpG within any cell-type. The second step involved a marginal conditional model that tested for evidence of differential DNAm within a particular cell type, adjusted for other cell types at a CpG. The joint model can be considered as an ANOVA test and provides evidence of differential DNAm in at least one cell type. The p-values were subjected to adjustment for multiple tests using the Benjamini-Hochberg.

We first estimated cell type proportion for each sample. The cell type proportion values from the whole blood and PBMCs were estimated with Houseman method using GLINT (ver. 1.0.4) script. Granule cell type was removed from the analyses because of low proportion of granule cells following PBMC isolation. Cell type proportion for each cohort is present in **Supplementary Figure 3**. TCA method was conducted following R TCA package workflow description to produce the methylomes of all cell types based on cell type proportion generated in Houseman analysis.

After TCA-deconvolution for five cell types, we removed CpG sites with variation < 0.002 in each cohort. As shown in **Supplementary Figure 4**, top 30 principal component (PCs) derived from the methylation in bulk cells was strongly correlated with cell types prior to TCA-deconvolution. However, the correlation of top 30 PCs on methylation in individual cell types was significantly decreased, suggesting that TCA robustly deconvoluted methylation for individual cell type and removed cell type confronting effects.

#### ***Cell-type level epigenome-wide association analysis***

We applied EWAS methods previously described by Zhang et. al[11]. on the methylome of each cell type. Two-step general linear models (GLM) were performed. The residual value of each CpG site was derived from the GLM model 1 to adjust the biological confounders and calculate top 30 principal components (PCs) on residual matrix. We performed correlation analysis of top 30 PCs with cell type proportion. The second GLM model adjusted PCs on residuals to capture the global unknown confounders besides all known confounders in the GLM model (1).
